# A quick protocol for assessing the therapeutical effect of treatments against *Candidatus* Liberibacter asiaticus using rooted *Citrus medica* cuttings

**DOI:** 10.1101/2025.10.02.680075

**Authors:** Beatriz Cristina Pecoraro Sanches, Talita Alves dos Santos, Eduardo Silva Gorayeb, Nelson Arno Wulff, Franklin Behlau

**Affiliations:** Fund for Citrus Protection, Research and development Department, Araraquara, 14807-040, Brazil

## Abstract

Huanglongbing (HLB), caused by *Candidatus* Liberibacter asiaticus (*C*Las), is the most devastating citrus disease worldwide. Developing effective therapies remains a major challenge, as *C*Las cannot be cultured *in vitro* and colonizes the host phloem systemically. This study presents a rapid, reproducible, and cost-effective *in vivo* platform for screening bacteriostatic and bactericidal compounds using *C*Las-infected citron (*Citrus medica* (L.) Osbeck) stem cuttings. Among seven citrus genotypes tested, citron stem cuttings exhibited superior rooting performance and uniform vegetative growth. Four propagation protocols were developed and assessed based on the dynamics of rooting and shoot growth, *C*Las colonization, and the response to oxytetracycline (OTC) treatment. *C*Las+ stem cuttings were treated with OTC via drenching at different developmental root stages. The roots or new vegetative flushes were sampled for bacterial quantification by qPCR. In Protocol 2, in which treatments were applied and sampled 14 and 35 days after planting (DAP) respectively, OTC-treated roots achieved the highest suppression of *C*Las and the lower incidence of *C*Las+ rooted cuttings compared to non-treated roots. Time-course analysis showed that OTC delayed bacterial establishment in root tissues, with maximal suppression observed at 35 DAP. The proposed protocols simulate the natural progression of systemic infection in citrus plants, allow the assessment of phytotoxicity, and offer a scalable technology that does not underestimate the efficacy of future bactericidal candidates. This platform significantly reduces time and cost compared to traditional seedling or nursery tree experiments and enhances the early-phase screening of antimicrobial compounds. Altogether, this stem cutting-based approach represents a biologically relevant, scalable tool to accelerate therapeutic discovery and strengthen integrated HLB management strategies.

## Introduction

Huanglongbing (HLB) is currently the most devastating citrus disease globally, causing severe economic losses across the Americas, Asia, and Africa^1^. The disease not only reduces yield and accelerates premature fruit drop^2^ but also results in smaller and more acidic fruits on symptomatic branches^3^. The primary pathogen associated with HLB is the α-Proteobacteria *Candidatus* Liberibacter asiaticus (*C*Las), which is vectored by the Asian citrus psyllid (*Diaphorina citri* Kuwayama, Hemiptera: Psyllidae)^4^. In plants, *C*Las is a phloem-limited bacterium inoculated into the plant during psyllid feeding^5^, which rapidly migrates to developing root and leaf tissues^6^. The phloem colonization by bacterial proliferation impairs the transport of nutrients and triggers an overproduction of reactive oxygen species (ROS) by the host, exacerbating tissue damage and chlorosis symptoms^7^.

HLB has been the major contributor to the sharp decline in citrus production in Florida, where output has fallen from 150 million boxes in 2005, when the disease was first detected in the USA, to just 18 million boxes in 2024^8^. Since 2013, 100% of Florida’s citrus orchards have been infected with *C*Las^9^. In Brazil, HLB was first detected in 2004 and is currently established in almost all key citrus-growing regions. The implementation of a comprehensive management program, which includes the use of healthy nursery trees, scouting and removal of infected trees, and a strict schedule of insecticide sprays, significantly slowed the disease progress in Brazil. Despite ongoing efforts, the incidence of HLB-infected trees in the São Paulo state citrus belt has risen significantly, from 24% in 2022 to 47% in 2025^10,11^. This increase is attributed mainly to the development of psyllid resistance to insecticides belonging to the classes of pyrethroids, neonicotinoids, and organophosphates in several important regions of São Paulo state combined with the maintenance of infected plants in the field^12^. During the 2024/2025 season, HLB was responsible for 9.05% of premature fruit drop^11^. In this context, there is growing interest in exploring novel therapeutic strategies, such as trunk injection of antibiotics, biotechnological approaches, and the prospection of new substances, for suppressing bacterial titer within affected plants.

Numerous attempts have been made to culture or co-culture *C*Las in laboratory settings^13–16^, but none have achieved successful results. This difficulty suggests that *C*Las has a strict metabolic dependence on the phloem sieve tubes, psyllid cells, and hemolymph for its physiological and reproductive needs^17^. One approach for developing treatments against *C*Las involves the use of *Liberibacter crescens* (*Lcr*) strain *BT-1*, the only cultured wild-type strain of the *Liberibacter* genus, which has been instrumental in some studies^18,19^. However, genomic differences between the two bacteria, along with the non-pathogenic nature of *Lcr*, limit its usefulness as a model organism for *in vitro* assays. Another tool for screening bactericidal compounds is the hairy root assay^20,21^, which utilizes *Agrobacterium rhizogenes* to induce transgenic expression of HLB-resistant candidate genes or to test antimicrobial activity in roots already pre-colonized with *C*Las. Since the tissue is infected prior to antibiotic treatment, this model does not replicate the natural progression of HLB, where systemic bacterial colonization occurs gradually in whole-plant systems. This limitation could potentially lead to an underestimation of an antibiotic’s efficacy, as the model doesn’t account for factors like bacterial movement and host responses that occur during early infection stages in mature citrus trees.

Vegetative propagation through stem cuttings is one of the most widely employed methods for producing herbaceous and woody plants globally^22^. The method allows for maintaining genetic uniformity, as the stem fragments will generate clones of the mother plant. However, only plants that possess a high capacity for adventitious rooting, rapid callus formation, and tolerance to water stress are eligible to be reproduced by this method^23^. In the context of the HLB pathosystem, the propagation of infected plants through stem cuttings can be advantageous, since the bacterium is systemic, and newly infected plants are rapidly generated. As living plant material, *Citrus spp.* stem cuttings enable the natural translocation of bacteria from source to sink, closely replicating the behavior observed in seedlings and mature orange trees in the field. Infected *Citrus* spp. stem cuttings could therefore serve as an *in vivo* model for screening novel bactericidal agents against HLB. Hence, *C*Las growth and translocation to newly developed tissues can be assessed following treatment with candidate substances. Furthermore, stem cuttings develop roots, produce new shoots, and reach maturity more quickly than seed-grown plants, enabling faster experimental outcomes. Compared to tissue culture or grafted seedlings, stem cuttings are often more cost-effective and can be propagated without the need for specialized equipment.

This study presents protocols based on *C*Las-infected citron stem cuttings to rapidly and cost-effectively assess the therapeutic potential of candidate compounds. This approach effectively mimics the HLB pathosystem, allows for the assessment of phytotoxicity across different substances, and facilitates the early identification of (bio)chemical treatments capable of suppressing *C*Las within plant tissues.

## Materials and methods

### Production of *Citrus* spp. stem cuttings

One-year-old citron plants, grafted onto rangpur lime (*Citrus × limonia* Osbeck) rootstock, were inoculated with *C*Las via grafting^24^. All plants were grown in 300-mL conical tubes (6.5 × 5.9 cm, upper × lower diameter; 16 cm, height), filled with coconut fiber, and irrigated twice a week. Six months after grafting, mature lateral flushes were collected from these *C*Las-infected *C. medica* potted plants. Stem cuttings measuring 6–8 cm in length and 4–5 mm in basal diameter were prepared, each with three nodes and a single leaf per node, which was trimmed to one-third of its original size to reduce moisture loss (Fig. 1). Immediately after cutting, the basal end of the stems was bevel-cut and immersed for 1 minute in a 7.5% (w/v) aqueous solution of indole-3-butyric acid (IBA; Sigma Aldrich, CAS No. 133-32-4). Stem cuttings were planted in 220 mL disposable plastic cups (95 mm height × 72 mm rim diameter), half-filled with autoclaved fine construction-grade sand (particle size 0.15–0.42 mm; 40–100 mesh). During protocol assessments, plastic cups were maintained at 26 °C under a 14 h light/10 h dark photoperiod, using a metal halide lamp as the light source (Philips HPI-T, 1000 W/543 E40) positioned ∼1 m above the plants. Watering was performed daily to form a minimal film on the sand surface (∼8 mL), ensuring consistent moisture. No fertigation was applied throughout the process.

**Figure 1.**
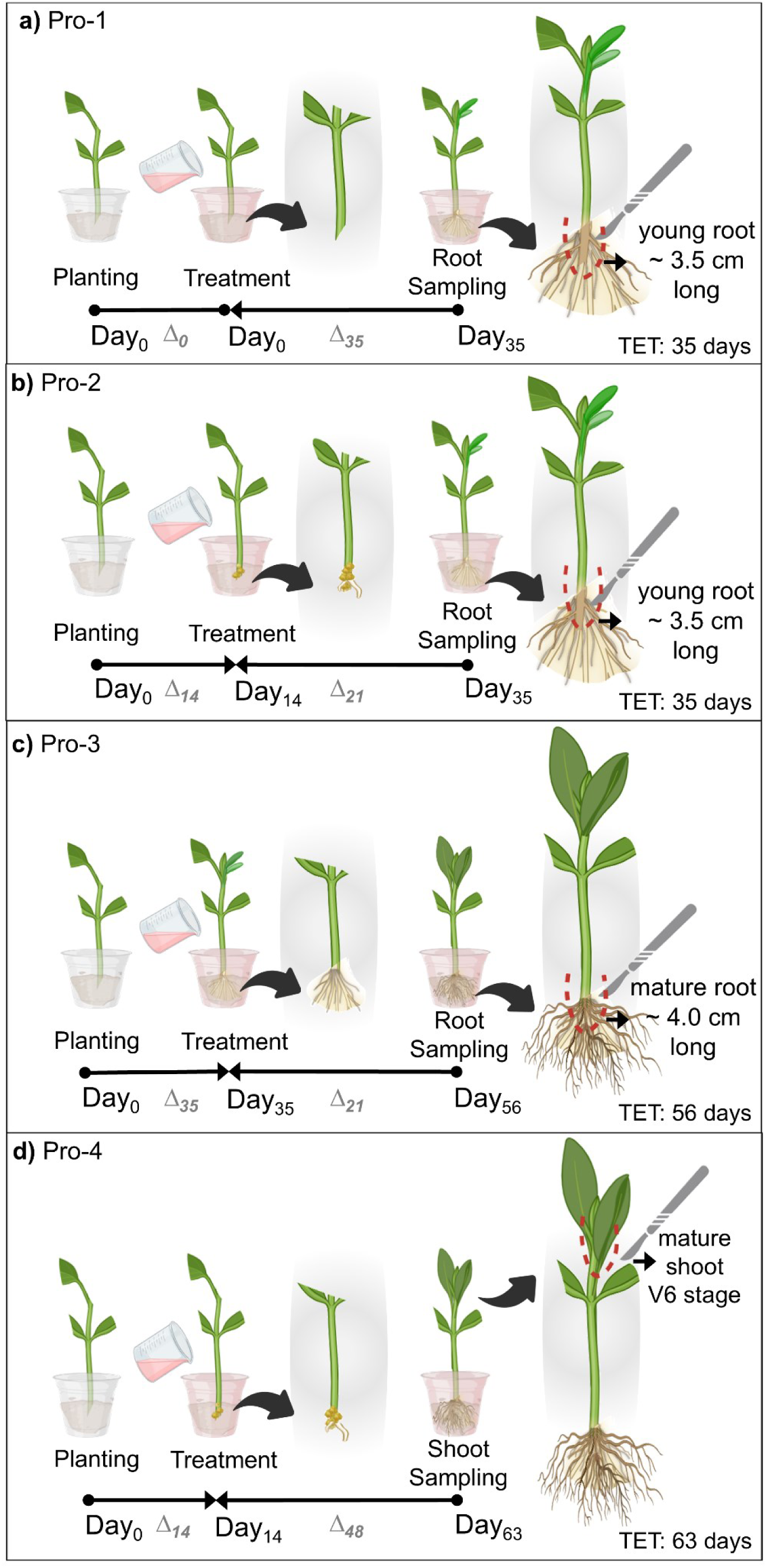
Schematic representation of the four protocols assessed for screening treatments against *Candidatus* Liberibacter asiaticus (*C*Las)-infected citron (*Citrus medica*) stem cuttings. (**a**) Pro-1: Stem cuttings were planted and treated on the same day, followed by root sampling 35 days after treatment for *C*Las titer analyses. (**b**) Pro-2: Stem cuttings were treated 14 days after planting (DAP), at the callus initiation stage, followed by root sampling 21 days after treatment for *C*Las titer analyses. (**c**) Pro-3: Stem cuttings were treated 35 days after planting, after root development, followed by mature root sampling 21 days after treatment for *C*Las titer analyses. (**d**) Pro-4: stem cuttings were treated 14 DAP and shoots at V6 stage^25^ were sampled 48 days after treatment for *C*Las titer analysis. Black arrows indicate a close-up of root development at treatment and sampling. Δn indicates the interval, in days, between steps. TET, total elapsed time. The figure was created using BioRender (https://biorender.com/).

### Comparison of stem cutting propagation efficiency and root–shoot development across citrus genotypes

Stem cuttings of seven citrus genotypes were compared regarding root and shoot development and propagation viability: citron (*Citrus medica*), alemow (*Citrus macrophylla* Wester); Pera, Valencia, Hamlin sweet oranges, (*Citrus × sinensis* (L.) Osbeck); Fortune mandarin (*Citrus reticulata* Blanco), and Persian lime (*Citrus latifolia* (Yu. Tanaka) Tanaka). Fifteen healthy stem cuttings were produced for each genotype, following the methodology previously described, and cultivated for nine weeks. Stem cutting viability (%), defined as the percentage of stem cuttings that successfully developed, root growth, and ontogeny of new flush were assessed weekly. New flush ontogeny was determined according to Cifuentes-Arenas et al (2018)^25^. Root development was assessed by measuring the length of the three longest roots of each cutting. Two independent experiments were conducted.

### Protocols for screening therapeutic treatments against *C*Las using rooted citron stem cuttings

Four distinct protocols using rooted stem cuttings of citron were assessed for their effectiveness in screening potential treatments against *C*Las (Pro-1 to 4, Fig. 1). Each experiment consisted of two treatments, treated and non-treated stem cuttings, with 15 stem cuttings (replicates) per treatment. Treated stem cuttings received oxytetracycline hydrochloride (OTC-HCl, Sigma-Aldrich, CAS N°: *2058-46-0*), used as a positive control. This molecule is the most effective antibiotic against bacteria and has been used in trunk injection therapies in the field^26^. Water-treated stem cuttings were used as negative controls. For all protocols evaluated, stem cuttings were obtained from infected *C. medica* plants exhibiting *C*L*as* titers of log₁₀ ≥ 6.8 (Ct ≤ 22.0). A single soil drench application using 10 mL of OTC at 0.625 mg/mL was performed in each planted cutting for all experiments. To minimize light-induced degradation of OTC, all treated stem cuttings were maintained in the dark for 24h immediately after OTC application. The plants returned to the previously established growth conditions after this period. All protocols were assessed in three independent experiments, with forty-five stem cutting samples for each protocol (n = 45). The main steps in each protocol was carried out as follows: Pro-1: OTC was applied to the sand immediately after planting the stem cuttings, followed by total root sampling 35 days after planting (DAP), when the roots had reached 3.0–3.5 cm in length; Pro-2: OTC was applied at 14 days after planting (DAP), coinciding with the onset of basal tissue regeneration and visible callus, characterized by the organization of a discrete epidermal layer and the formation of a root cap^27^; root sampling was performed at 35 DAP; Pro-3: OTC was applied 35 DAP, followed by total root sampling at 56 DAP, when roots had reached ∼4.0 cm length; Pro-4: OTC was applied 14 DAP, but instead of roots, new flushes at the V6 stage^25^ were sampled at 63 DAP. Root and shoot samples were submitted to qPCR analyses to measure *C*Las titer.

### DNA extraction and qPCR analysis

The complete root system (Pro-1, Pro-2, and Pro-3) and the central leaf nerve of young shoots (Pro-4) from each cutting were collected and chopped with sterilized blades. Tissue sample aliquots of ∼0.3 to 0.5 g were added to 2.0 mL Eppendorf microtubes and processed following the protocol of Teixeira et al. (2005)^28^ using a TissueLyser II system (Qiagen, Valencia, CA, USA) at 45 Hz for 30 seconds, with 5 mm steel beads. Total DNA was extracted using the cetyltrimethylammonium bromide (CTAB) method described by Murray and Thompson, 1980^29^.

Real-time PCR (qPCR) for the detection of *C*Las was performed using 1.0 μL of total DNA (100 ng/μL), TaqMan® PCR Master Mix (5x) (Applied Biosystems), and HLBas primers/probe at final concentrations of 0.4 μmol/L and 0.2 μmol/L, respectively. Reactions were carried out in a StepOnePlus™ thermocycler (Applied Biosystems, California, USA) in a total volume of 12 μL^30^. The thermal cycling conditions were as follows: 50 °C for 2 minutes, 95 °C for 20 seconds, followed by 40 cycles of 95 °C for 3 seconds and 60 °C for 30 seconds^30^. *C*Las quantification was based on the linear relationship between Ct values and the logarithmic concentration of *nrdB* rRNA, using the transformation log_10_(10^−0,2921∗Ct value+11,62^) ∗ 66,67 ÷ 5. Samples were considered *C*Las-positive when the qPCR cycle threshold (Ct) was below 35.0^30,31^. Notably, based on the qPCR analysis criteria, only samples with *C*Las+ titers, i.e., above 2.5 log₁₀ cells/g of tissue (Ct ≤ 35.0), were included in the statistical analyses and graphical representations.

### Time course of *C*Las titer in roots of OTC and non-treated stem cuttings

The population dynamics of *C*Las in infected stem cuttings were monitored to identify the optimal sampling moment, defined as the timing at which *C*Las concentration in roots between OTC and non-treated plants differed significantly. Stem cuttings from *C*Las-infected citron plants were prepared and treated with OTC, following the Pro-2 methodology. Water-treated stem cuttings were used as negative controls. Complete root system of each cutting was sampled weekly over 77 days after planting (DAP). DNA extraction and *C*Las titer quantification were determined as previously described.

### Data analyses

The homogeneity of variances and normality of the data were assessed using Bartlett’s and Shapiro–Wilk tests, respectively. Data on cutting viability, *C*Las population titers, and the dynamics of *C*La*s* population growth in OTC and non-treated stem cuttings were compared using Student’s *t*-test (α = 0.05). The incidence of *C*Las+ stem cuttings was compared using the non-parametric Wilcoxon Mann Whitney test. All statistical analyses were performed in R software (v. 4.5.0)^32^.

## Results

### Comparison of citrus genotypes for cutting propagation

Citron displayed one of the highest rooting efficiencies among the tested genotypes, showing both rapid development of new roots and shoots and high viability from stem cuttings. Cutting viability, e.g., the number of rooted and well-established stem cuttings by the total number of planted cuttings, were, in order, 90.0%, 86.7%, 86.2%, 80.0%, 70.0%, 43.3% and 33.3%, respectively, for alemow, ‘Valencia’, citron, Persian lime, ‘Pera’, ‘Hamlin’ and mandarin (Fig. 2a). Citron was the first to initiate root development from stem cuttings at 14 DAP, achieving an average root length of 30.0 mm by 35 DAP, when root elongation stabilized. Alemow and Persian lime initiated rooting at 21 DAP and reached average root lengths of 25.0 mm and 11.3 mm by 35 DAP, respectively. While root growth of alemow halted at 35 DAP, roots of Persian lime continued developing until 42 DAP. The sweet orange cultivars ‘Pera’, ‘Valencia’, and ‘Hamlin’, initiated root development later, around 28 DAP, reaching an average root length of ∼10 mm by 56 DAP. Mandarin was the last genotype to initiate root development, at 35 DAP, and reached a maximum length of ∼ 5 mm by 56 DAP (Fig. 2b). A similar pattern was observed for flush development (Fig. 2c). Only genotypes that established a root system earlier showed consistent vegetative growth within the assessed period. Citron progressed more rapidly and uniformly to the V6 stage than alemow, with 66.7% stem cuttings at V6 and 33.3% at V5 by 63 DAP, versus 26.6% of cuttings at V4, 46.7% at V5, and 20.0% at V6 in the same period, respectively. In contrast, ‘Pera’ had only 14.3% of the cuttings at V4, whereas ‘Valencia’, ‘Hamlin’, mandarin, and Persian lime did not flush up to 63 DAP.

**Figure 2.**
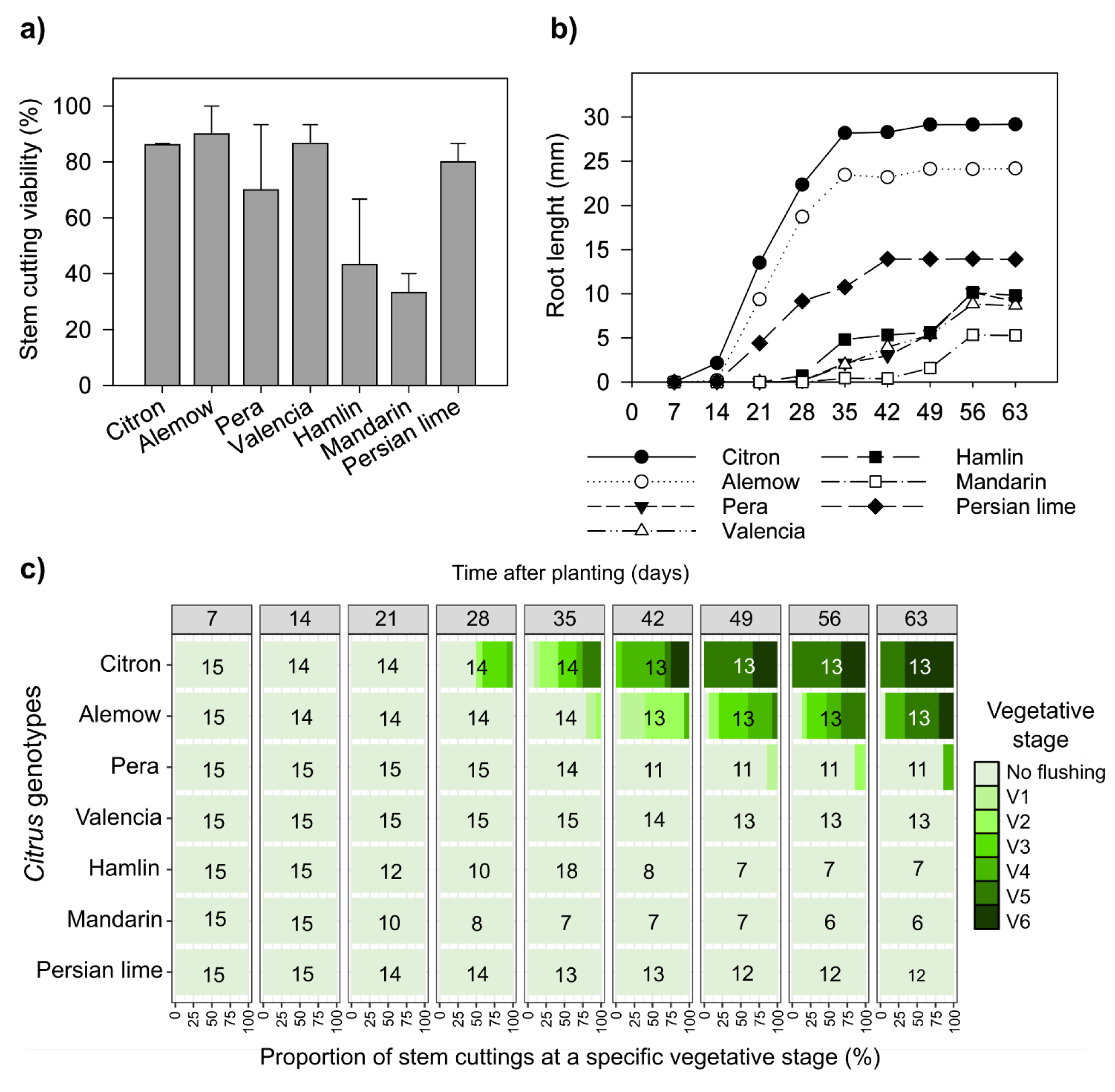
Viability, root development, and vegetative flush ontogeny of citron (*Citrus medica*), alemow (*C. macrophylla*), Pera, Valencia and Hamlin sweet oranges (*Citrus × sinensis*), Fortune mandarin (*C. reticulata*), and Persian lime (*C. latifolia*). (**a**) Percentage of viable stem cuttings at 63 days after planting (DAP), means ± standard errors of mean from two independent experiments (n = 25–30). (**b**) Development curve depicting weekly root length development (mm) up to 63 days after planting (DAP), as means from two independent experiments (n=25–30). (**c**) Presence of vegetative flush (%) ontogeny stages assessed weekly up to 63 DAP classified according to Cifuentes-Arenas et al. (2018)^25^, as means of one experiment. Numbers inside the square boxes indicate the number of viable replicates over time (n).

### Therapeutic treatment protocols

Cutting viability was similar between OTC-treated and non-treated stems across all protocols assessed (Fig. 4a, c, e, g). Survival rates were 55.6% in OTC-treated versus 66.7% in non-treated cuttings in Pro-1 (*t* = −1.06; *p* = 0.30; Fig. 4a), 66.7% versus 64.7% in Pro-2 (*t* = 0.25; *p* = 0.81; Fig. 4c), 80.0% versus 77.8% in Pro-3 (*t* = 0.25; *p* = 0.81; Fig. 4e), and 73.4% versus 77.8%, in Pro-4 (*t* = −0.31; *p* = 0.76; Fig. g).

**Figure 3.**
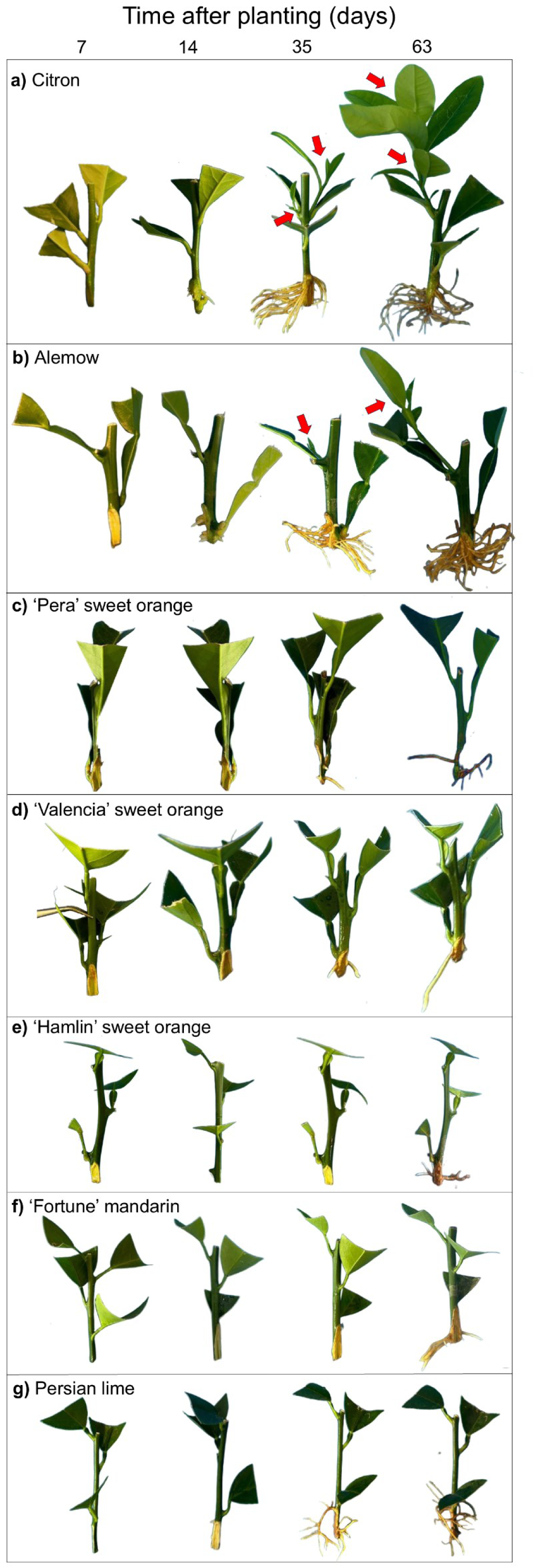
Ontogeny of root development and vegetative flush at 7, 14, 35 and 63 days after planting of (**a**) citron (*Citrus medica*), (**b**) alemow (*C. macrophylla*), (**c**) Pera, (**d**) Valencia, and (**e**) Hamlin sweet orange (*Citrus × sinensis*), (**f**) Fortune mandarin (*C. reticulata*), and (**g**) Persian lime (*C. latifolia*). Red arrows indicate new vegetative flushes. All stem cuttings measured 7 to 8 cm long at planting.

**Figure 4.**
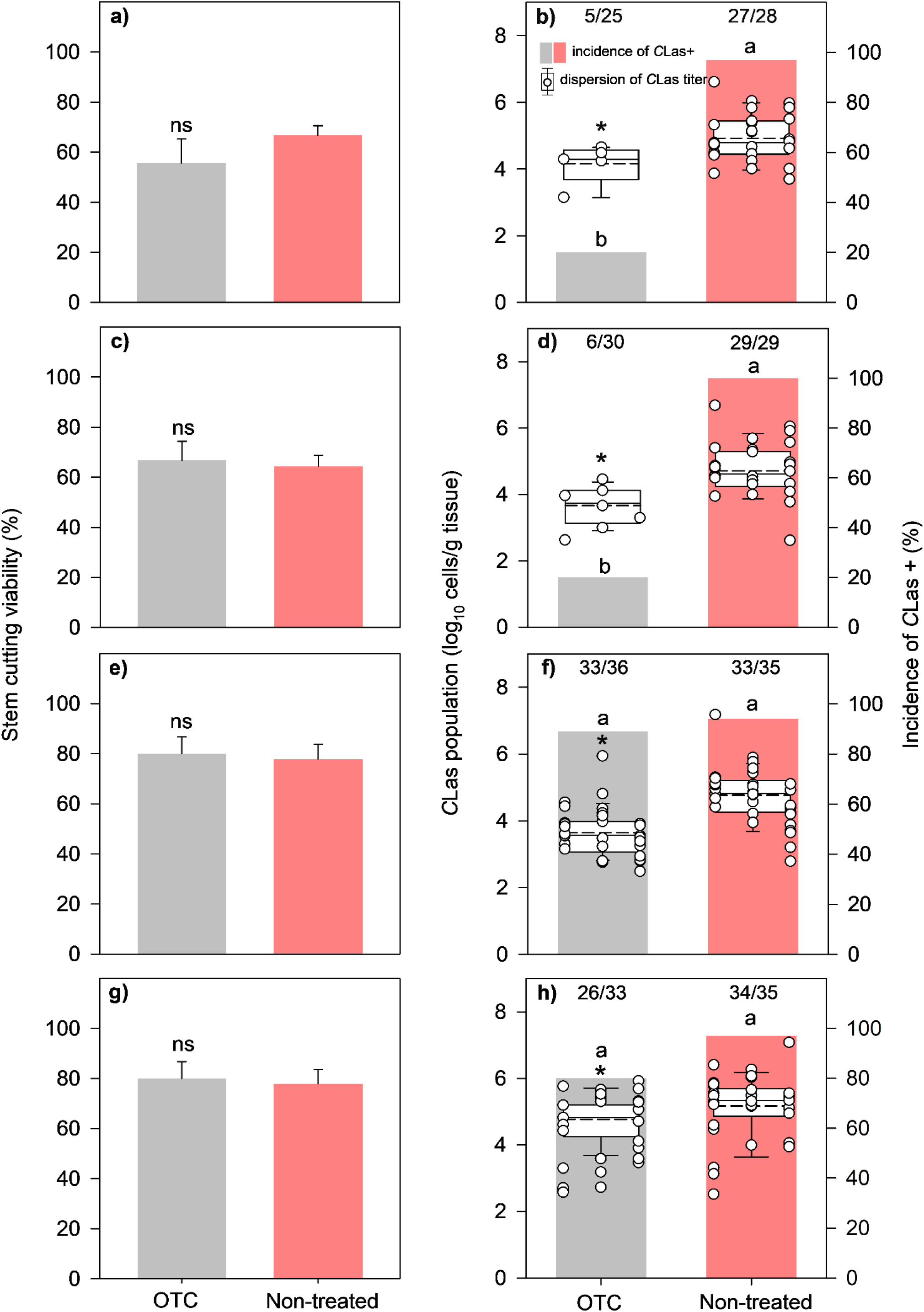
Viability of citron (*Citrus medica*) stem cuttings (%) (**a**, **c**, **e**, **g**), *C*Las incidence (%), and population (log_10_ cells/g of tissue) (**b, d, f, h**), in roots or shoots, collected after treatment. Panels represent the four experimental protocols: (**a**–**b**) Pro-1, (**c**–**d**) Pro-2, (**e**–**f**) Pro-3, and (**g**–**h**) Pro-4. In **a**, **c**, **e**, **g**, data are presented as the mean of three independent assays ± standard error of the mean. ns indicates no significant difference between treatments (Student’s *t-* test, α = 0.05). In **b**, **d**, **f**, **h**, the box plot displays the distribution of bacterial titers in log₁₀ (cells/g tissue) for three separate assays, with solid and dashed lines indicating the mean and the median values, respectively. Stacked white circles correspond to independent experiments. Stem cuttings that tested *C*Las-(log₁₀ ≤ 2.5, Ct ≥ 35.0) or yielded undetermined titers were excluded from box-plot analysis. * indicates significant difference in log₁₀ means between treated and nontreated stems (Student’s *t*-test, α = 0.05). Different letters above columns indicate significant differences in the incidence of *C*Las+ stems between treated and nontreated stems (Wilcoxon rank-sum test, α = 0.05). The number of CLas+ stems over the total number of viable stems used is shown above the bars.

The proportion of *C*Las+ cuttings varied markedly among protocols (Fig. 4b, d, f, h). In Pro-1 and Pro-2, OTC treatment substantially reduced *C*Las incidence. Only 20.0% of treated cuttings tested positive in both protocols, compared to 96.7% and 100.0% in non-treated groups in Pro-1 and Pro-2, respectively (*p* < 0.01; Figs. 4b and 4d). In Pro-3 and Pro-4, the proportion of *C*Las+ cuttings was similar between treatments. In Pro-3, 91.7% of the OTC-treated cuttings were *C*Las+ as opposed to 94.3% in non-treated cuttings (*p* = 0.67; Fig. 4f). In Pro-4, the proportion of *C*Las+ in treated and non-treated stems was 78.8% and 97.2%, respectively (*p* = 0.02; Fig. 4h).

Bacterial population in roots and leaves was reduced after OTC treatment in all protocols (Fig. 4b, d, f, h). In Pro-1, OTC-treated root cuttings averaged a lower *C*Las population of 4.1 log₁₀ cells/g, compared to 4.9 log₁₀ cells/g in non-treated cuttings (t = –2.21; p < 0.05; Fig. 4b). The OTC effect was more pronounced in Pro-2, in which titers declined to 3.3 log₁₀ cells/g under OTC treatment as opposed to 4.7 log₁₀ cells/g in untreated cuttings (t = –4.57; p < 0.01; Fig. 4d). A similar pattern was observed in Pro-3, with OTC-treated roots averaging 3.7 log₁₀ cells/g compared with 4.7 log₁₀ cells/g in non-treated (t = – 5.70; p < 0.01; Fig. 4f). In Pro-4, *C*Las titer in new leaves reached 4.4 log₁₀ cells/g after OTC treatment as opposed to 5.2 log₁₀ cells/g (t = –2.98; p < 0.01; Fig. 4h) in untreated cuttings.

### Temporal dynamics of *C*Las multiplication in roots

As Pro-2 (Fig. 5 a-c) was the most suitable protocol for comparing treated and untreated stems, an experiment was carried out to assess the temporal dynamics of *C*Las multiplication following this protocol. OTC-treated stem cuttings significantly modified the early colonization dynamics of *C*Las in newly formed citrus roots, delaying bacterial establishment (Fig. 6a-b). At 7 and 14 days after planting (DAP), *C*Las detection did not reach the detection threshold in OTC-treated and non-treated roots in both experiments. At 28 DAP, the bacterial population of the non-treated stem cuttings grew exponentially, reaching 4.6 and 4.9 *C*Las log₁₀ cells/g in the first and second replicates, respectively.

**Figure 5.**
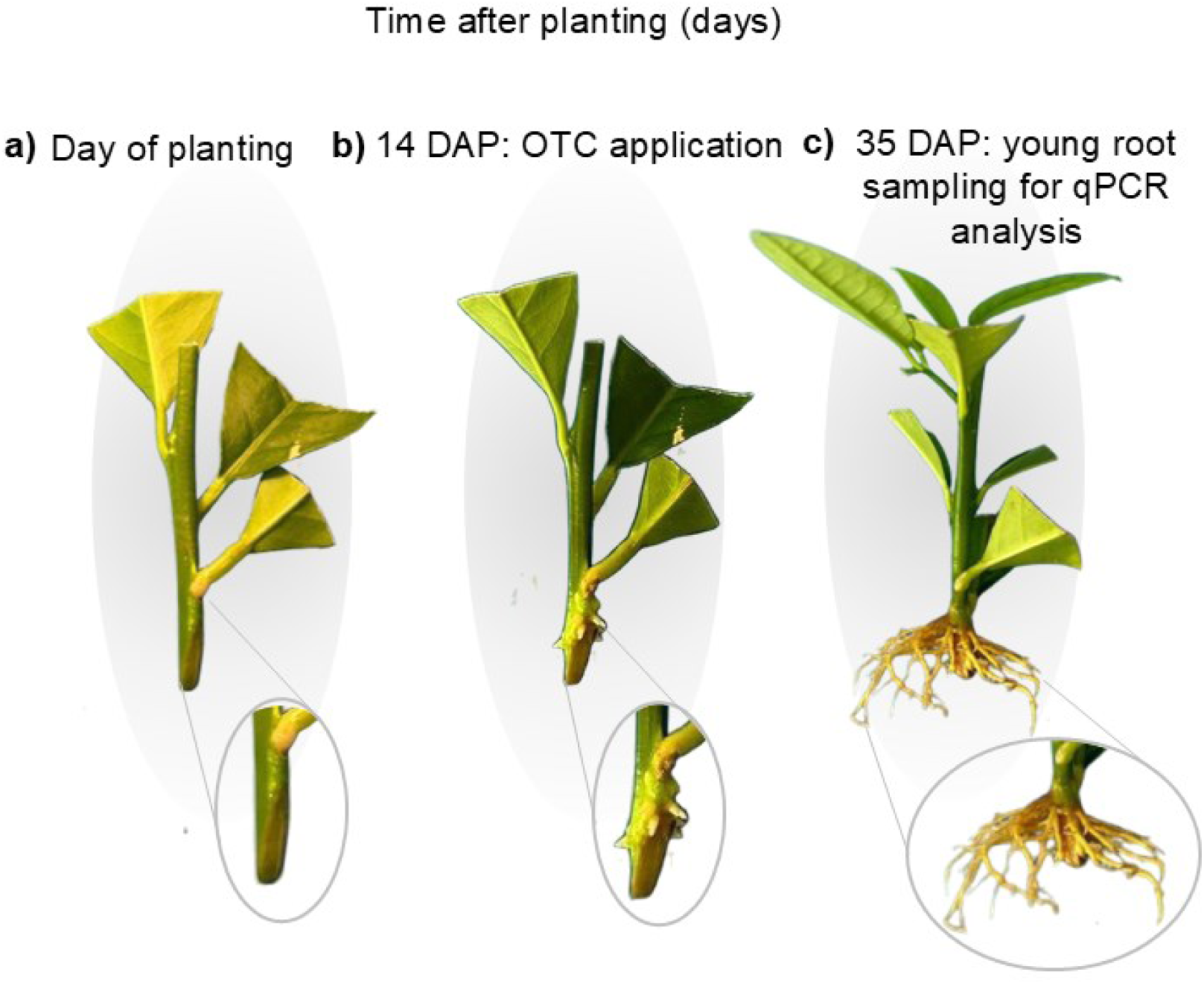
Root development stages of *Citrus medica* stem cuttings following Pro-2 methodology, with a close-up of the three steps, (**a**) stem cuttings at day of planting, (**b**) callus formation and initial root emergence at 14 days after planting (DAP) and (**c**) established root system at 35 DAP, when roots were sampled for qPCR analysis.

**Figure 6.**
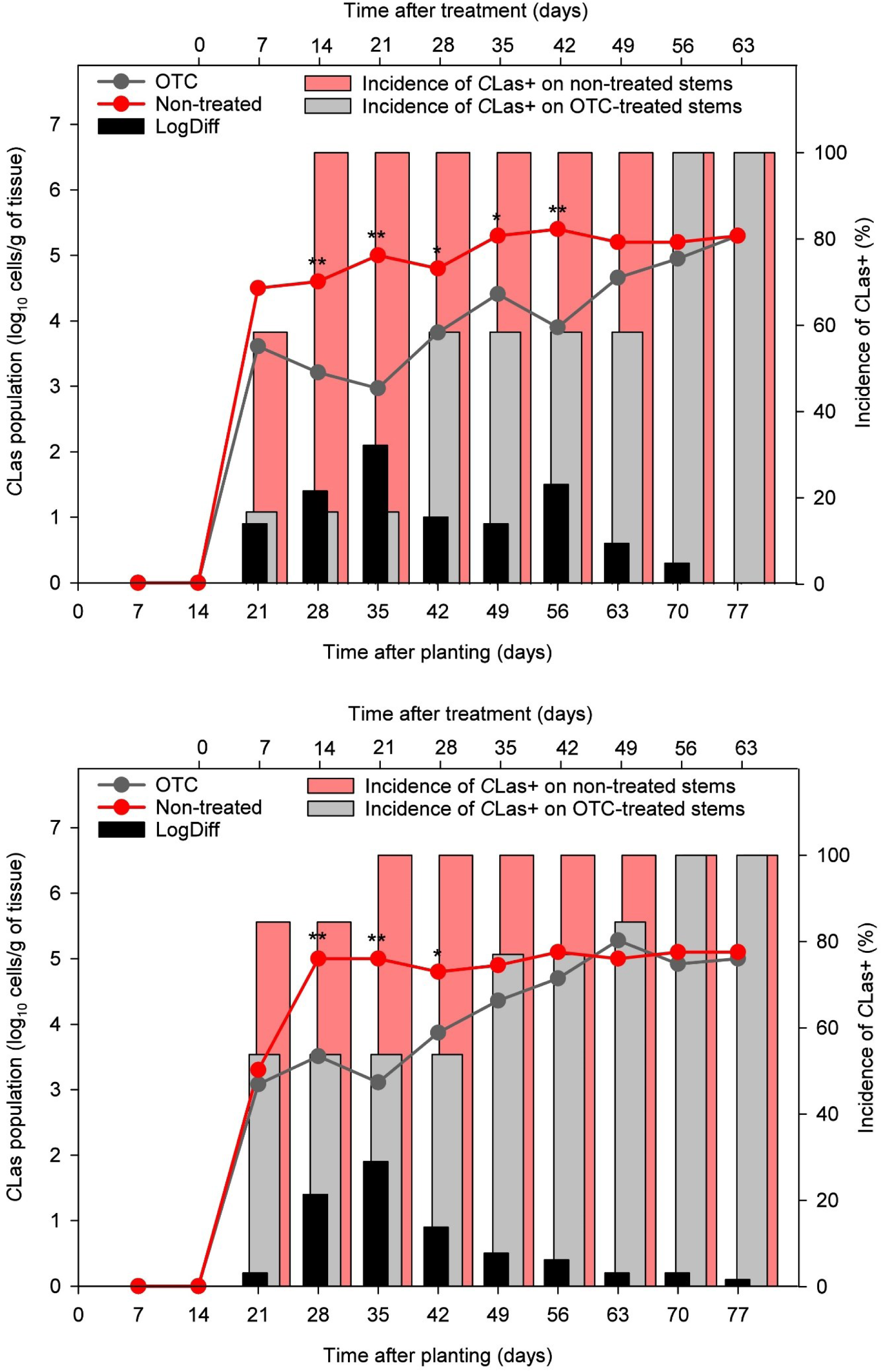
Dynamics of *Candidatus* Liberibacter asiaticus (*C*Las) population in newly formed roots of *Citrus medica* cuttings treated or non-treated with oxytetracycline (OTC) over 77 days after planting, in biological replicates (**a**) 1 and (**b**) 2. LogDiff, logarithmic difference between OTC and non-treated rooted cutting stems. Data are shown as means ± standard errors of the mean (n = 13). Stem cuttings that tested *C*Las-(log₁₀ ≤ 2.5, Ct ≥ 35.0) or yielded undetermined titers were excluded from the analysis. * indicates significant difference in Log₁₀ means between treated and nontreated stems (Student’s *t*-test, \**p* ≤ 0.05, \*\**p* ≤ 0.01).

The rapid increase, followed by stabilization over time, indicates successful colonization and a sustained bacterial presence in the roots of untreated stems throughout the 77-day assessment period. In contrast, OTC-treated roots exhibited significant lower *C*Las population at 28 (*t* = −3.60, *p* < 0.01), 35 (*t* = −4.85, *p* < 0.01), 42 (*t* = −2.34, *p* < 0.03), 49 (*t* = −2.66, *p* = 0.02) and 56 DAP (*t* = −6.59, *p* = 0.01) in the first replication, and at 28 (*t* = −9.36, *p* < 0.01), 35 (*t* = −8.55, *p* < 0.01) and 42 (*t* = −2.33, *p* = 0.03) in the second replication. By 63 DAP, *C*Las titers in treated roots converged with those observed in the non-treated cuttings, reaching, respectively, 4.6 versus 5.2 *C*Las log₁₀ cells/g (*t* = - 1.79, *p* = 0.08) in the first replication and 5.3 versus 5.0 *C*Las log₁₀ cells/g (*t =* 0.77, *p =* 0.44) in the second repetition. The lowest *C*Las titer in roots of OTC-treated stems was observed at 35 DAP, averaging 3.0 and 3.1 *C*Las log₁₀ cells/g, in the first and second assays, respectively. During this period, the greatest numerical difference in *C*Las titer between roots of treated and untreated stems was observed, reaching 1.9 and 2.1 log₁₀ *C*Las cells/g in the first and second replicates, respectively.

## Discussion

In the present study, a quick and easily accessible protocol was developed to optimize the screening of antimicrobial molecules against *C*Las. Currently, there are no effective or widely adopted therapeutic treatments for HLB. Given the increasing incidence of HLB-infected trees in prominent citrus growing areas, a dramatic rise in the psyllid population due to insecticide resistance, and the substantial economic loss caused by the disease, there has been a growing interest in therapies targeting *C*Las in infected trees^3,33,34^. In this context, this study meets the urgent need for the development of a reliable *in planta* methodology to rapidly assess the efficacy of candidate compounds or strategies aimed at suppressing the bacterium in infected citrus organs. Moreover, the proposed protocol overcomes the limitation that *C*Las remains unculturable under standard, axenic laboratory conditions, which significantly constrains experimental approaches for screening and validation of potential treatments^16^.

The new methodology accounts for characteristics of the selected host. Citron was selected as the model plant due to its early, uniform, and high ability of stem cuttings to generate roots^35^, vigorous shoot development, and susceptibility to *C*Las colonization. Using a genotype that roots more easily is preferable, not only to shorten the duration of the assays but also to enhance the absorption of the treatments being evaluated. The successful uptake and systemic movement of antimicrobial compounds in citrus are dependent on the physicochemical properties of the product and the physiological traits of the host, particularly new flushes and root development, which vary among citrus genotypes. Previous studies have shown that citron, lemon, and lime root more readily than mandarins and sweet oranges^36–38^. Likewise, Ghorbel and collaborators^39^ reported that citron and alemow showed superior horticultural performance compared to other assessed citrus genotypes, with rooting efficiencies of 85.0%. By contrast, ‘Pera’ sweet orange showed a low rooting efficiency^40^, consistent with other *C. sinensis* cultivars studied in the present work. Similarly, although Persian lime and mandarin had some root development, it was slower than that observed for citron and their stems failed to produce new vegetative flushes within the assessment period, limiting the suitability of these genotypes for rapid screening applications. Overall, our results strengthen the variability in rooting capacity among citrus species and highlight citron as the most suitable genotype for the proposed protocol.

Proof-of-concept studies employing biotechnological approaches and trunk injection of antibiotics in nursery trees of sweet orange have demonstrated promise in suppressing *C*Las infection, providing a direct method for delivering bactericidal agents into the citrus vascular system^26,41^. These approaches effectively circumvent limitations associated with foliar applications and the restricted systemic movement of antimicrobial agents^42,43^. Despite the diversity of compounds tested, only OTC applied through trunk injection has shown limited effect in treating HLB-affected trees in the field^26^, underscoring the need for searching alternative bactericides. OTC does not cure the disease or eliminate *C*Las from the tree, but reduces the bacterial titer inside the plant^44^. This could potentially minimize psyllid transmission and help mitigate disease damage, allowing partial recovery of affected trees and extending the productive lifespan of infected plants^45^. Although antimicrobial peptides, nanoparticles, and small-molecule antibiotics have also demonstrated potential results in reducing *C*Las titers in HLB-diseased trees^21,46–48^, none of these technologies has been proven consistently effective or has been widely adopted in the field. Despite some advances, the high-throughput screening of new molecules utilizing trunk injection systems remains labor-intensive, time-consuming, and demanding innovative methodologies^17,49,50^. For instance, considering all the steps, a single assessment based on trunk injections of compounds into previously HLB-infected citrus seedlings conditioned in greenhouse conditions takes at least one year to complete^46,47^. By contrast, the protocol developed in this study enables a preliminary screening of candidate compounds in just 35 days, offering a significantly faster and more scalable alternative.

Four protocols were assessed, considering the critical biological aspects of the pathosystem, including uneven distribution of the bacterium within host organs, systemic translocation, and colonization dynamics toward new vegetative flush and young root systems^6,51,52^. The stem-cutting approach favors increased contact between the applied compound and the plant vascular tissues, enhancing absorption via the xylem and potentially promoting its radial translocation into the phloem through apoplastic loading^46,53,54^. Moreover, as roots continue to elongate, the process naturally generates microinjuries in the epidermis and cortex, which may facilitate compound penetration and further increase uptake efficiency^55,56^. This dynamics promotes a more effective delivery of the antimicrobial agent to the phloem, where *C*Las resides, thereby increasing the likelihood of direct interaction between the compound and the pathogen.

In protocol Pro-1, in which stems were drenched immediately after planting, and Pro-2, in which treatment was drenched two weeks after roots began to emerge, stem cuttings that received OTC showed a reduction in *C*Las titers and a higher proportion of *C*Las-negative samples compared to the respective untreated controls. These findings suggest that both protocols offer suitable conditions for detecting antimicrobial activity and can serve as effective platforms for identifying promising bactericidal candidates, which further supports the use of such screening strategies. However, given the unknown effects of a candidate compound on cutting viability and potential phytotoxicity, Pro-2 may be the preferred option, as it allows initial root development before treatment. In contrast, Pro-1 remains a viable alternative for treatments that do not hinder or may even stimulate rooting. Notably, both protocols contribute meaningfully to early-stage screening strategies, offering flexibility depending on the physicochemical and biological properties of the candidate compounds under assessment.

By contrast, Pro-3 resulted in a lower reduction of bacterial titer after treatment; however, no difference in the incidence of *C*Las+ was observed between OTC-treated and non-treated roots. This is possibly related to the timing of drench application, as *C*Las was already well established in the root system at the time of treatment initiation (35 DAP). Using pre-colonized roots may lead to an underestimation of a treatment’s therapeutic effect. Similar challenges have been reported in other studies using pre-infected tissues, as in assays with cysteine-rich peptides in excised infected leaves^57^ and treatments with ferulic acid and bioflavonoids in hairy root systems^21^.These methodologies showed limited efficacy even for OTC, a therapy currently recognized as effective in reducing bacterial titer. Likewise, in Pro-4, which was based on sampling young shoot tissue of drenched stems, bacterial titers from OTC-treated stems were lower than titers from non-treated stems; however, no significant difference in the incidence of *C*Las+ stems was found when new flushes from OTC-treated and non-treated cuttings were sampled, suggesting that this protocol may underestimate the therapeutic potential of a candidate treatment. Additionally, Pro-4 is more time-consuming, as it requires an extended period for shoot emergence and leaf sampling. Taken together, these findings reinforce Pro-2 as the most suitable and efficient protocol for screening antimicrobial treatments against *C*Las.

The temporal dynamics of root colonization observed in this study support the use of 35 DAP as a reliable sampling point for *C*Las detection, as implemented in Pro-2. The most pronounced differences in bacterial populations between OTC-treated and nontreated stems occurred at 28, 35, and 42 DAP, with log₁₀ reductions of 1.4, 1.9, and 0.9 in the first experiment and 1.4, 2.1, and 1.0 in the second experiment, respectively. Before 21 DAP, *C*Las root colonization was absent or incipient, with inconsistent differences between experiments and treatments. At later stages, 49 and 56 DAP, *C*Las titers remained lower in OTC-drenched compared to untreated stem cuttings, although reductions were less pronounced than those observed between 28 and 42 DAP. Conversely, by 63 DAP, no difference was observed between treatments, suggesting a systemic recolonization of the root system. These findings indicate that the root system represents an early and favorable environment for *C*Las proliferation and detection, corroborating previous studies that identified roots as primary sites of systemic infection^6,58^.

The proposed methodology offers the advantage of assessing bactericidal efficacy directly against *C*Las within citrus tissues under controlled and reproducible conditions. Using citron as a *C*Las propagation system provides several logistical and experimental benefits, as a single plant can yield multiple uniform stem cuttings that facilitate replication, reduce costs, and significantly accelerate the screening process. Moreover, this method enables early detection of phytotoxic effects, which is critical for assessing the practical feasibility of field applications and mitigating potential physiological damage to the host^43,59^. As a result, this system allows for more accurate and efficient evaluation of antibacterial activity. Rather than replacing traditional nursery plant-based assessments, this platform serves as a practical and biologically relevant first-tier screening tool for the rigorous and cost-effective pre-selection of promising antimicrobial candidates. Collectively, these features make the presented protocol a robust and scalable strategy for early-stage antimicrobial screening against HLB.

## ACKNOWLEDGEMENTS

The authors thank Elaine C. Martins for sharing the quantification curve data for *C*Las titer estimation, and Isabela V. Primiano for the contribution on Figure 2.

## Data availability

The manuscript and supporting files include all data generated during this study.

## Author contributions

B.C.P.S. and F.B. designed research. E.S.G. analyzed data. B.C.P.S., T.A.S. performed research. N.A.W. provided *C. medica* plants and reagents. B.C.P.S wrote the manuscript. All authors reviewed and approved the final version of the manuscript.

